# Cardinal v3 - a versatile open source software for mass spectrometry imaging analysis

**DOI:** 10.1101/2023.02.20.529280

**Authors:** Kylie Ariel Bemis, Melanie Christine Föll, Dan Guo, Sai Srikanth Lakkimsetty, Olga Vitek

**Affiliations:** Khoury College of Computer Sciences, Northeastern University, Boston, USA; Institute of Surgical Pathology, Medical Center – University of Freiburg, Faculty of Medicine, Freiburg, Germany; German Cancer Consortium (DKTK) and German Cancer Research Center (DKFZ), Heidelberg, Germany

## Abstract

Cardinal v3 is an open source software for reproducible analysis of mass spectrometry imaging experiments. A major update from its previous versions, Cardinal v3 supports most mass spectrometry imaging workflows. Its analytical capabilities include advanced data processing such as mass re-calibration, advanced statistical analyses such as single-ion segmentation and rough annotation-based classification, and memory-efficient analyses of large-scale multi-tissue experiments.

## Main

Mass spectrometry imaging (MSI) provides unique value for life science research. MSI analyzes spatial distributions of hundreds of analytes directly from complex biological samples such as tissue sections at cellular resolution. Typical analytes include lipids, metabolites, drugs, peptides and proteins. The untargeted, label-free and multiplexed measurement capabilities make MSI an up-and-coming technology for research applications in medicine, biology, pharmacology, toxicology and forensics^1^.

The nature of MSI data challenges its analysis. A single MSI file often reaches dozens of gigabyte in size. MSI data from a single tissue section comprises several thousand mass spectra, each containing thousands of mass-to-charge (m/z) feature – intensity pairs. Each spectrum is annotated with x- and y-coordinates that localize it on the tissue section. Further spectra annotations such as type of tissue, disease or treatment are often stored in a separate file. However, a typical MSI experiment is even more complex, and consists of multiple tissue sections and conditions. This data complexity and size requires specialized and efficient software tools for data preparation, preprocessing and statistics, which differ from classical mass spectrometry experiments.

Many vendor-independent, free and open source MSI software exist as recently reviewed by Weiskirchen and colleagues^2^. A commonly used MSI software is the commercial SCiLS Lab software (Bruker Daltonics, Bremen,Germany). Non-commercial MSI software is either i) freeware such as DataCubeExplorer^3^ and MsIQuant^4^; ii) open source software build on a proprietary programming language that are based on costly licenses (MATLAB) such as MSiReader^5^ and SpectralAnalysis^6^; or iii) open source software with permissive-license such as rMSI^7^, BASIS^8^, M^2^aia^9^, HIT-MAP^10^ and massPix^11^.

While permissive-license open source software is a prerequisite for reproducible research, the respective MSI software are often specialized towards specific analysis tasks. A typical MSI analysis requires multiple analysis steps: data import, visualization, preparation, preprocessing, statistics & artificial intelligence (AI), and data export. These are linearly depicted in Fig. 1a, but are rather iterative in practice. Most open-source software target particular steps in this analytical pipeline. MassPix focuses on lipid analysis and identification, while HIT-MAP is specialized towards peptide analysis and identification. BASIS focuses on preprocessing methods. M2aia offers methods for multi-modal imaging analysis. rMSI suite consists of several different R packages, which together offer solid support for most analysis steps (data import, preparation, preprocessing, visualization and identification), but only offers basic capabilities for downstream analyses. Beyond specific functionalities, the large-scale nature of the MSI data makes it necessary to deploy these tools on specialized hardware, such as cloud computing. The targeted scope of the existing open-source tools, and the requirements for a computational infrastructure in most cases, limit their usability for many labs.

**Figure 1.**
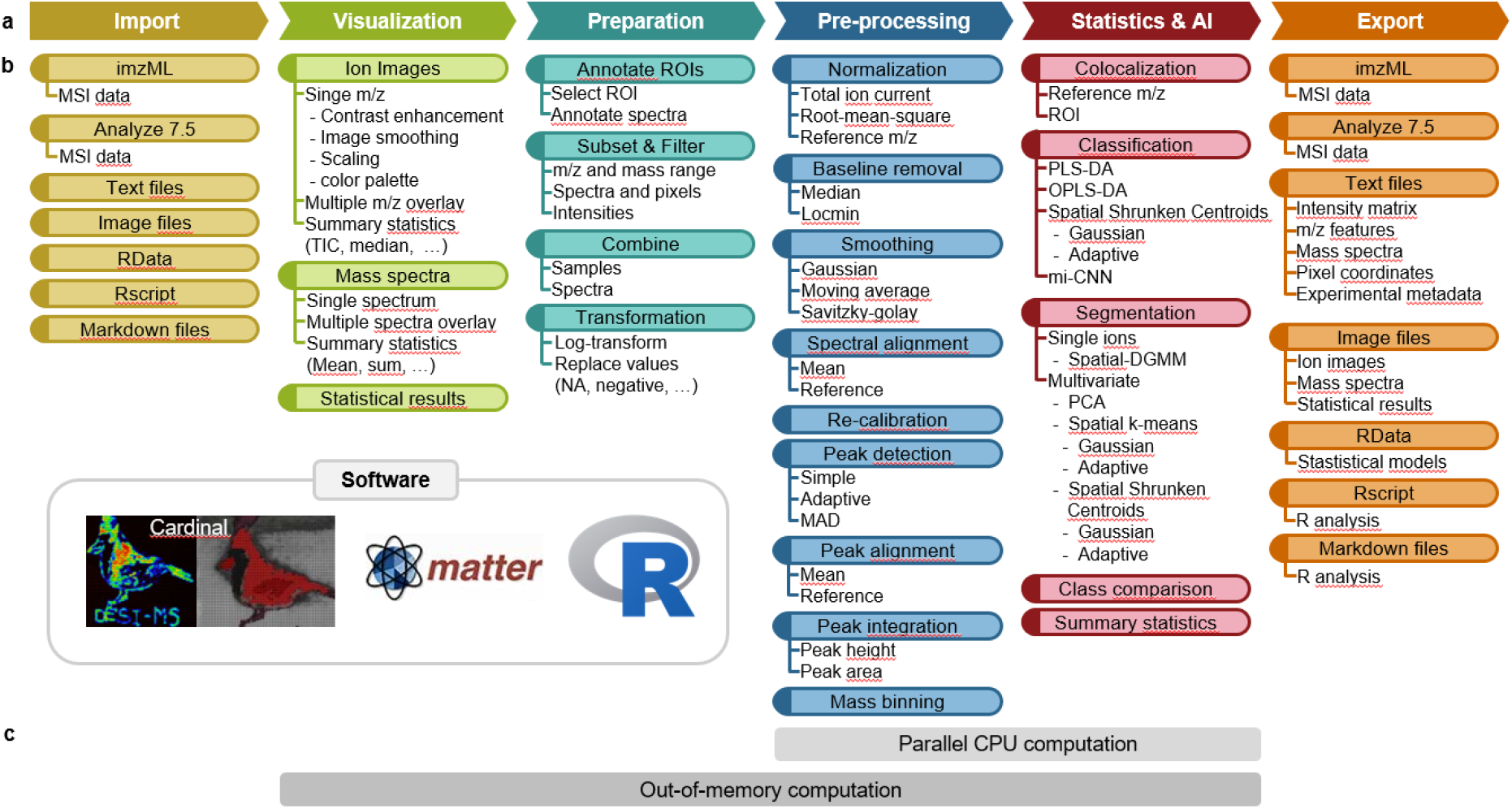
Common MSI data analysis steps and the corresponding functionalities in Cardinal. **a**, MSI data analysis include import, visualization, preparation, preprocessing, statistical analysis & artificial intelligence (AI), and export. **b**, Preprocessing methods in Cardinal allow adjustment of the analysis of various types of MSI data. Statistical and AI methods enable class discovery (image segmentation), class comparison (detection of changes in abundance) and class prediction (tissue and spectra classification). **c**, Cardinal supports the analysis of large files on traditional hardware via parallel and out-of-memory computing. Time efficient parallel CPU computation is enabled for all preprocessing and some statistics functions via the R package BiocParallel. Most Cardinal functions directly interact with larger than memory files via the Matter R package, without the need for converting data or copying files. **Abbreviations:** DGMM: Dirichlet Gaussian mixture models; MAD: mean absolute deviations; OPLS-DA: orthogonal partial least squares discriminant analysis; PCA: principal component analysis; PLS-DA: partial least squares discriminant analysis; ROI: region of interest; TIC: total ion current, mi-CNN: multiple instance convolutional neural network.

Unlike the methods above, Cardinal v3 is a permissive-license family of open source R/Bioconductor packages. It has one of the richest portfolios of advanced statistical and machine learning methods (Fig. 1), as well as the capability to work with large-scale data in a standard desktop computational environment.

A recent study^12^ highlighted Cardinal’s versatility by reviewing its deployment for different analytes (metabolites, lipids, peptides and proteins), different ionization sources (matrix-assisted laser desorption/ionization (MALDI), desorption electrospray ionization, secondary-ion mass spectrometry), and mass analyzers (time-of-flight (TOF), orbitrap). In the same study^12^, Cardinal and the commercial SCiLS Lab software were applied for the analysis of *Pseudomonas aeruginosa* colonies. Both software produced comparable results for unsupervised segmentation and finding discriminative m/z features. Thus, Cardinal represents not only one of the most comprehensive open-source MSI software but also a competitive one.

Compared to Cardinal v1^13^ (and the unpublished Cardinal v2), Cardinal v3 was improved in three different aspects. i) Existing methods for MSI data import and export, preprocessing, and visualization were refined, now allowing for example the analysis of high mass resolution data, spectral mass alignment and re-calibration; ii) a broad scope of statistical and machine learning functionalities was added to find a) analytes with the same distribution by co-localization analysis; b) differentially abundant analytes via class comparison^14^; c) analytes with homogeneous and heterogeneous spatial distributions by single-ion segmentation^15^; d) deep learning based methods for weakly supervised classification were developed^16^ in python and implemented as CardinalNN R package to not burden Cardinal users to learn another programming language; iii) major restructuring of the underlying code infrastructure supports the efficient analysis of large experiments via out-of-memory computation and parallel computation.

In the following, we illustrate how Cardinal v3 enables new, and principally different research. We highlight four reproducible case studies based on open datasets from the PRIDE repository^17^: 1) Segmentation of a high resolution phospholipid imaging data set^18^, 2) Single ion segmentation and concentration curves of a very large sized peptide imaging data set^19^, 3) Supervised and 4) Semi-supervised classification of a multiple replicate peptide imaging data set^20^. The raw data were transferred to the MassIVE database and the R Markdown files containing the R code for the case studies were added as re-analysis via MassIVE.quant (MassIVE identifier: MSV000086099, MSV000086102, MSV000089594).

The first case study demonstrates Cardinal v3 capabilities to accurately and efficiently handle high mass resolution datasets, which was not possible with Cardinal v1. We preprocessed the mouse bladder phospholipid imaging data and then reproduced figures from the original publication (Fig. S1). Furthermore, we applied spatial shrunken centroids (SSC), Cardinal’s unique spatially aware unsupervised segmentation method^21^. Unsupervised SSC resulted in five spatial segments corresponding to the three different bladder tissue types and two background segments (Fig. 2a). SSC performs feature regularization to report segment specific m/z features. These contained not only m/z features, which were described in the original publication as tissue specific, but also many additional ones (Fig. 2b).

**Figure 2.**
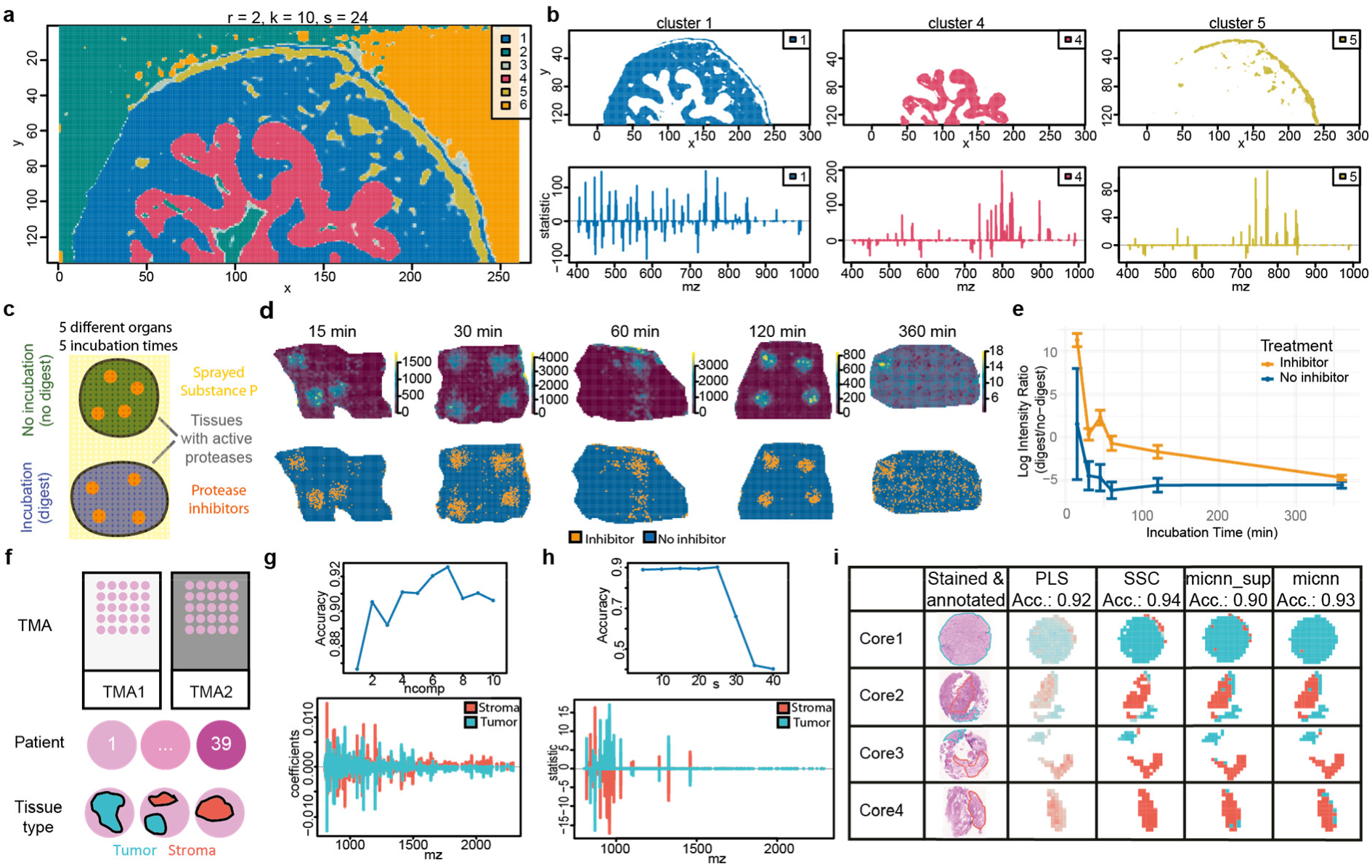
Re-analysis of three public datasets with Cardinal. Three MSI experiments were chosen as case studies: 1) a high-resolution segmentation dataset (a-b), 2) a large-scale single ion dataset (c-e) and 3) a multiple replicates classification study (f-h). **a**, Case study 1: spatially aware unsupervised segmentation of mouse bladder tissue produced five spatial segments, corresponding to muscle layer (blue), adventitial layer (yellow), urothelium (magenta), background (green) and an unknown cluster (orange). **b**, Each tissue segment shows a distinct statistical contribution profile of its m/z features. **c**, Case study 2: experimental design. Fresh-frozen porcine tissues from five different organs were covered with substance P as a protease substrate tracer. Substance P was either applied before or after incubation (digested vs. non-digested condition) for five different durations. Four spots with protease inhibitors were added to each tissue section to prevent digestion in these areas. **d**, Ion image of substance P (m/z 1347.5) and segmentation of substance P at five different incubation times. Single ion segmentation with two segments finds segments corresponding to the areas with (orange) and without (blue) protease inhibitor mix. **e**, For both segments, the log intensity ratio for mean substance P in digested and inhibitor region at all five incubation times was plotted to show the tissue protease activity with different incubation times. **f**, Case study 3 and 4: experimental setup. Two tissue micro arrays (TMA) contained 39 patient’s bladder tissue cores with tumor and stroma annotations. **g**, Cross validation reveals the best parameters for partial least squares(PLS). The contribution of the different m/z features to the two classification categories, tumor and stroma, are plotted below. **h** Best parameters and contribution of m/z features to classification for spatial shrunken centroids (SSC). **i**, Class annotation annotated by a pathologist in the stained tissues next to spectra-wise class prediction by PLS, SSC, multiple instance convolutional neural network with (micnn_sup) and without detailed spectra annotations (micnn) for four exemplary tissue cores. Classification accuracy (Acc.) for each method on the complete test dataset is stated.

The second case study showcases Cardinal v3’s ability to analyze large datasets with highly customized and flexible analysis steps including the new and unique single-ion segmentation method. The analyzed 55 GB dataset is one of the largest public MSI datasets and represents 78 fresh-frozen porcine tissue sections from five organs, and contains regions with different protease activity due to different digestion times (Fig. 2c). Calculating the mean of all 281,395 spectra took ∼ 1,200 seconds and ∼ 9 GB memory on a personal computer. Parallel computing with two cores reduced calculation times to ∼ 700 seconds. Further preprocessing took ∼ 1000 seconds and ∼ 10 GB memory. Thus, compared to Cardinal v1 and most non-commercial MSI software, Cardinal v3 enables the analysis of larger than memory datasets in reasonable times without the need to invest too much into computational infrastructure. Cardinal v3’s unique single-ion segmentation method revealed the spectra with and without protease activity (Fig. 2d), which were not publicly available. Basic R functionalities enabled calculating and plotting the protease activities in different segments and at different time points (Fig. 2e). This highlights the sheer unlimited options of the R environment, which goes far beyond the customization options that stand-alone MSI software can provide.

The third case study benefits from Cardinal v3’s new preprocessing and highlights the performance of Cardinal’s unique SSC method for tissue classification and m/z feature selection for complex experimental designs. The dataset contains two files with 39 patient bladder tumor tissue cores and is accompanied by spectra annotation such as tissue type (Fig. 2f). In Cardinal, we directly attached these spectral metadata to the raw data to minimize mistakes. We applied the new mass alignment and mass re-calibration preprocessing methods, which successfully reduced mass shifts and improved mass accuracy, which is key for correct analyte matching and identification (Fig. S2 a,b). One file was used for training and cross validation to find optimal parameters for classification, either with the spatial shrunken centroids or the partial least squares method. Both methods are tailored towards classification of spatially dependent data and their optimal parameters were found by cross validation (Fig. 2g). Spatial shrunken centroids includes only the most discriminating m/z features into the classifier, which facilitates m/z feature extraction and interpretation of the classifier (Fig. 2g). Both classifiers showed an accuracy above 90% to distinguish tumor and stroma spectra on the second file, which served as independent test set (Fig. 2h). This shows that Cardinal even enables the reproducible and successful analysis of lower quality datasets from experiments with multiple tissues and conditions. The fourth case study illustrates Cardinal’s new deep learning extension for semi-supervised classifications. The new method performs sub-tissue classifications with only tissue-level annotations using multiple instance learning and captures m/z dependencies using convolutional neural networks. The same multiple replicate dataset as in the third case study was used. The results show that this classifier achieved comparable accuracy on the test file compared to the classifier trained with sub-tissue level annotations (Figure 2h).

The distribution of the Cardinal software family as open-source R/Bioconductor packages provides an ecosystem that ensures software quality and maintenance through semi-annual releases, version control, detailed documentation and open development^22^. Additionally, Cardinal may be installed via Github, Bioconda^23^, and BioContainers^24^, or used on public clouds via the graphical user interface of Galaxy^25^ (Table 1). Cardinal’s open source license and its imzML export function facilitate data sharing, reproducibility, interoperability, and flexibility. Complete analyses done in Cardinal and shared via simple R scripts or markdown embedded code lay the foundation of reproducible research. In three published studies, Cardinal users adjusted and expanded functionalities to their specific needs and redistributed their new code or software for the benefit of the whole community^10,12,26^.

**Table 1:**
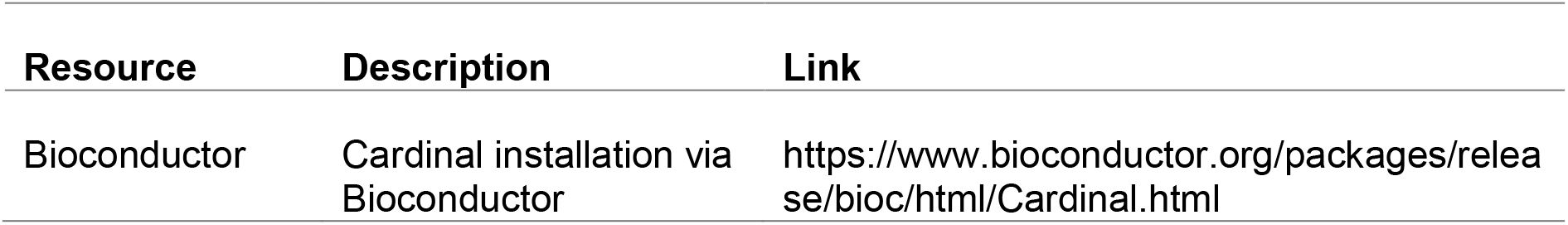
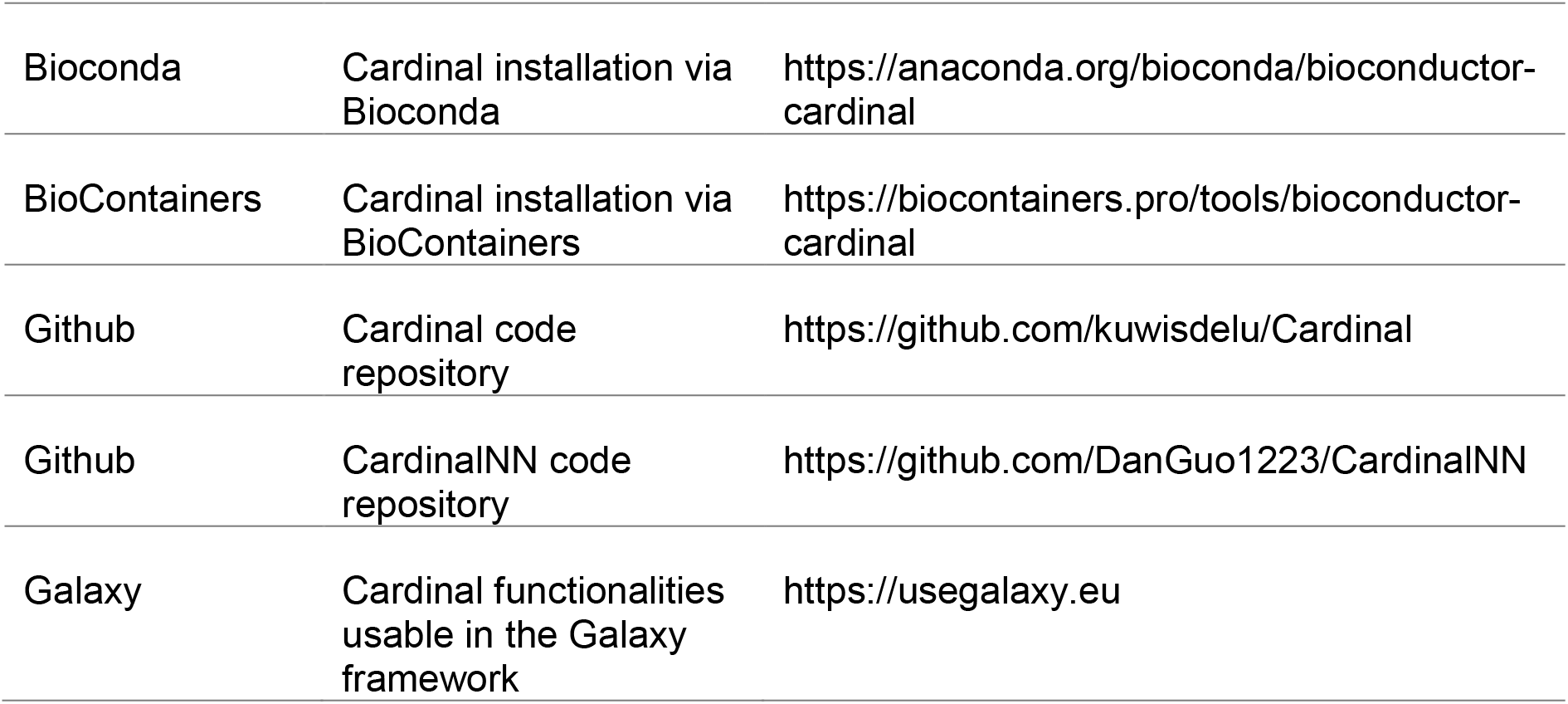
Cardinal is available via different resources

To maximize its usability, Cardinal provides extensive training and help infrastructure (Supplementary Table 1) including user guides, vignettes describing case studies and documentation of algorithms. Overall, the updated infrastructure and methods strengthen Cardinal v3 portfolio as a computationally efficient MSI software with a huge methods portfolio that supports the analysis of a diverse range of MSI experiments.

## Methods

### Case study 1: High resolution segmentation dataset

Römpp et al. imaged phospholipids in a mouse urinary bladder tissue section with an AP-SMALDI-LTQ Orbitrap mass spectrometer that allowed a mass resolving power of 30000 at m/z 400 and 10 µm spatial resolution^18^. The processed imzML file was downloaded from PRIDE (PXD001283) and imported into Cardinal using 10 ppm to obtain regular m/z bins. The average mass spectrum between m/z 770.4 and 770.7 was plotted to visualize two close peaks. Peak detection was performed on the mean spectrum with MAD noise estimation and a signal to noise ratio of six. Peaks were aligned to the mean spectrum using 15 ppm and only kept if their mean intensity was above zero. For each m/z in the generated peak list, the original peak area was integrated with the peakBin function in each total ion current normalized spectrum. After preprocessing, the overlaid ion image was plotted for the two close m/z features 770.51 and 770.56 using adaptive image smoothing, linear image normalization and contrast enhancement via suppression. The spatial distributions of three m/z features (741.5307, 798.5410, and 743.5482) were overlaid into one ion image plot using adaptive image smoothing, linear image normalization and contrast enhancement via suppression. Next, the preprocessed data was segmented using unsupervised spatial shrunken centroids^21^ using Gaussian weights with a radius (r) of two, maximum of ten clusters (k) and smoothing values (s) of 0, 6, 12, 18, 24 and 30. Overlaid and separated segment images as well as the statistics showing the contribution of each m/z feature were plotted for the segmentation with s = 24. A co-localization analysis was performed to find m/z features that correlate with m/z 743.5448, which is located in the lamina propria.

### Case study 2: Large scale single ion segmentation dataset

Erich et al. cut consecutive sections from five different porcine tissues (kidney, spleen, pancreas, liver and muscle) and spotted four times 1 µl protease inhibitor mix onto each tissue section^19^. For each organ, five tissue sections were spray-covered with substance P before incubation to allow its digestion in the areas without protease inhibitor mix (‘digested samples’) and the other six sections were spray-covered after incubation to prevent digestion (‘not digested samples’). Incubation times were set to 15, 30, 60, 120 and 360 min to measure the endogenous protease activity over time. Afterwards, imaging with a MALDI-TOF device in the m/z range 500 to 2500 at 200 µm spatial resolution was performed. The 55 GB imzML ‘time-curve-dataset’ was downloaded from PRIDE (PXD011104) and imported into Cardinal. Peak detection was performed on the mean spectrum with the MAD noise estimation and a signal to noise ratio of 2.5. Peaks were aligned to the mean spectrum and only m/z features with non-zero mean intensities were kept. The peak area was integrated in the total ion current normalized data at the m/z peak positions ± 100 ppm. Time and memory consumption for these calculations were recorded once while using a single core and once with two cores via the BiocParallel function. Next, the ion image of substance P (m/z 1347.7) was plotted on the preprocessed file with the following parameters: contrast enhancement via histogram and adaptive image smoothing. All tissue specimens were annotated for their condition (digest/no digest) and time points (15, 30, 45, 60, 120, 360 min) via their position in the x-y coordinate grid. The five different tissue types were manually annotated according to the annotation provided in the original manuscript. These spectra annotations were integrated with the MSI data and filtered for ‘digest’ and ‘no digest’ spleen spectra. Substance P (m/z 1347.7) was plotted in the ‘digest’ spleen tissues with contrast enhancement via suppression and Gaussian image smoothing. Single ion segmentation^15^ of substance P in the digested spleen was performed to obtain the spectra with and without inhibitor mix. This was performed by applying spatially-aware Dirichlet Gaussian mixture model (DGMM) with a radius (r) of one, two clusters (k) and without annealing during parameter optimization. The obtained spatial clusters were plotted. For both clusters, substance P mean intensity was calculated for the digested (no inhibitor mix) and not digestion (inhibitor mix) spectra of both datasets at all digestion times. The obtained mean intensities were normalized to the mean intensity of substance P in the undigested spleen tissues at the same time points, log transformed and plotted as a time curve.

### Case study 3: Multiple replicates classification dataset

Tryptic peptides were imaged in urothelial tissues with a MALDI-TOF device at 150 µm spatial resolution^20^. The study cohort consisted of two tissue microarrays (TMAs) containing 39 patient’s urothelial tissue specimens with different types of urothelial cancer or benign diagnoses and annotations for tumor and stroma tissue regions. Both Analyze 7.5 files as well as the metadata containing spectra annotations were downloaded from PRIDE (PXD026459) and imported into Cardinal. This metadata was attached to the raw data in Cardinal and the data was filtered to keep only spectra with tumor or stroma annotations. The preprocessing was done separately for TMA1 and TMA2 to borrow as little information as possible between the two datasets that serve as test and training dataset in the classification. Gaussian signal smoothing with a window of 8 and kernel standard deviation of 2 was performed. Next, the baseline was removed with a median function that was applied to 750 blocks. The m/z values of all spectra were aligned to their mean spectrum with 200 ppm tolerance. Zoomed in mass spectra for six random spectra before and after m/z alignment were plotted for visual control. To shift the m/z to the correct positions, m/z re-calibration with four internal calibrants and 200 ppm was performed. Again, six mass spectra before and after re-calibration were plotted, zooming onto the m/z axis for the internal calibrant angiotensin. Afterwards, peaks were detected in TMA2 with the ‘simple method’, signal to noise ratio of 5, window of 10 and 500 blocks to estimate the noise. Picked peaks were aligned with 200 ppm to the mean spectrum and filtered to keep only peaks that occur in at least 1% of all spectra and had a mean intensity above zero. The obtained peak list was used to integrate the corresponding peak intensity areas from the smoothed and baseline removed data in a 200 ppm window with the peakBin function in TMA1 and TMA2 separately. Lastly, intensities were normalized to the total ion current of each spectrum. Classification using spatial shrunken centroids algorithm^21^ was performed. First, to find the optimal classification parameters, 5-fold cross validation was performed on the training dataset (TMA2), while peak binning to the peak list was done separately for each fold. Next, the preprocessed TMA2 file was classified using spatial shrunken centroids classification with the optimal parameters (r = 1, s = 25). Then the classifier was used to predict the test dataset (TMA1). Classification (cross validation, classification and prediction) was repeated with the partial least squares (PLS) method.

### Case study 4: Multiple replicate semi-supervised classification

The processed dataset generated in case study 3 from the 39 urothelial tissue cores was used in this case study. The split of training and test set was according to the two TMAs in which the tissue cores were assembled. To mimic a scenario where spectra annotations are not available but only the annotation for the complete tissue are available, the tissue annotations were assigned as tumor if any spectrum in the tissue was tumor and as stroma if none of the spectra in the tissue was tumor. The training dataset with tissue annotations was used to train the semi-supervised classification model on one TMA. Note that the spectra annotations were not used during training. The classifier used was a convolutional neural network (CNN) with three convolutional layers and one fully connected layer^16^. Then, the trained CNN was used to predict spectra annotations on the testing dataset, the other TMA, and the metrics, such as accuracy, sensitivity, and specificity, were calculated based on the ground-truth spectra annotations. To compare with standard supervised training that uses the detailed spectra annotations, the CNN with the same architecture was trained using ground-truth spectra annotations and evaluated on the same test dataset.

## General methods

For all case studies, we uploaded the raw data together with the analysis R code as a re-analysis into the MassIVE.quant repository^27^ (MassIVE & MassIVE.quant identifier: case study 1: MSV000086099 & RMSV000000684; case study 2: MSV000086102 & RMSV000000664; case study 3 and 4: MSV000089594 & RMSV000000686) and in addition deposited the R code in GitHub (https://github.com/Vitek-Lab/Cardinal3-vignettes).

All analyses were performed with Cardinal (version 3.1.0) in R. Figures were exported from R as pdf file and figure as well as text size were adjusted in Adobe Illustrator.

## Acknowledgements

The authors would like to thank Andreas Weber for critically revising the manuscript and vignettes. This work was supported by awards NSF-BIO/DBI 1759736, NSF-BIO/DBI 1950412, NIH-NLM-R01 1R01LM013115 and Chan-Zuckerberg foundation to O.V. M.C.F. was supported by the Hans A. Krebs Medical-Scientist-Programme, Faculty of Medicine, University of Freiburg, Germany.

## Author contributions

K.A.B. developed the methods and wrote the code for the Cardinal and Matter software. D.G. developed the methods and wrote the code for the CardinalNN software. M.C.F. and S.S.L. tested all software. K.A.B., M.C.F. and D.G. developed and analyzed the data for the case studies and generated the vignettes. S.S.L. revised the vignettes. M.C.F. wrote the manuscript and generated the figures. M.C.F. and O.V. revised and edited the manuscript. O.V. supervised the work and provided funding.

## Supplementary Figures

**Supplementary Figure S1.**
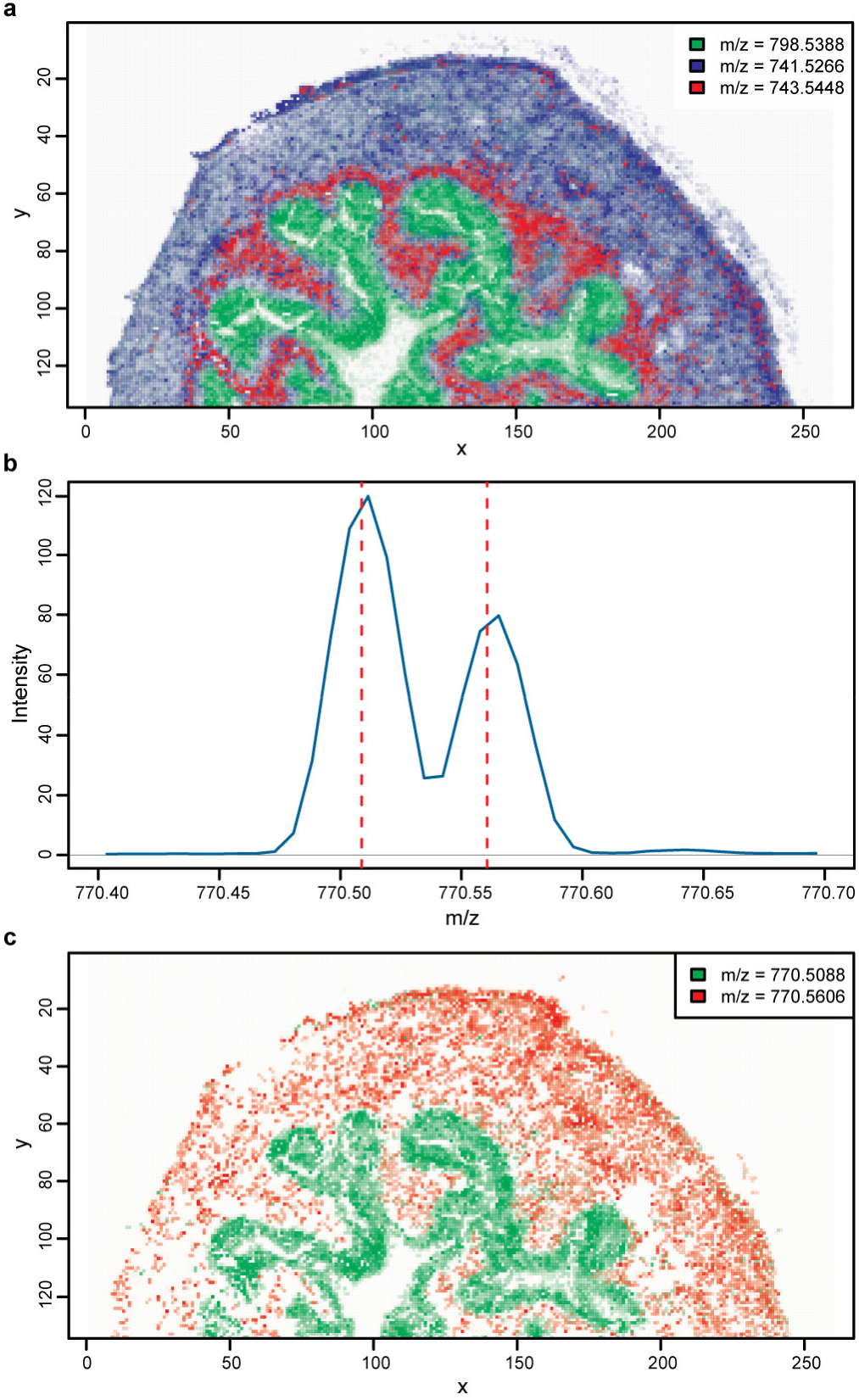
Cardinal 2 supports visualization and preprocessing of high mass resolution data. **a**, Reproducing the overlay ion image of three m/z features with different spatial abundances (Figure 1A in original publication). **b**, Zoomed in average mass spectra show two peaks that are only 0.05 m/z apart (Figure 2C in original publication) and could be detected as separate peaks. The red dotted lines indicate that both peaks will be picked separately during peak detection. **c**, Visualizing the spatial distribution of both peaks shows that they are present in different tissue regions (Figure 2B in original publication). The peaks in the original publication were likely mislabeled as suggested by Fig. S5 of the same publication and the inverse image we obtained here.

**Supplementary Figure S2.**
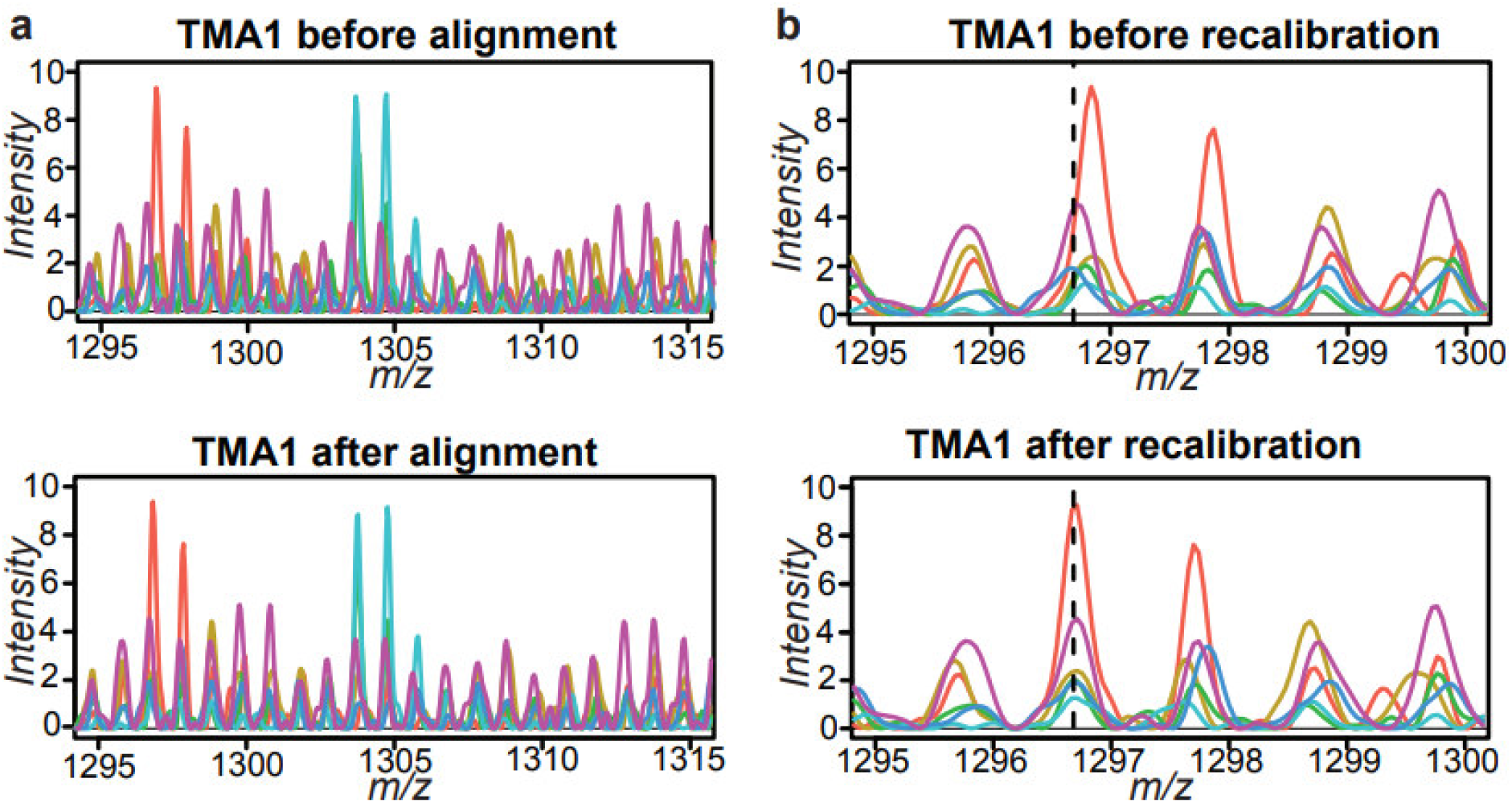
Cardinal 2 enables more accurate m/z values in the multiple replicates classification dataset. **a**, Zoomed in mass spectra for six random spectra of the first dataset show m/z shifts between the spectra before m/z alignment. Applying Cardinal’s new mass alignment step increases the alignment of the peaks substantially. **b**, Zoomed in mass spectra for six random spectra of the first dataset before and after mass re-calibration show how Cardinal’s new mass re-calibration method shifts the monoisotopic angiotensin peak towards its theoretical m/z position (m/z 1296.7, dashed vertical line).

## Supplementary information

Supplementary table 1: Cardinal documentation, training, support

